# Optimized detection of acute MHV68 infection with a reporter system identifies large peritoneal macrophages as a dominant target of primary infection

**DOI:** 10.1101/2020.11.17.387969

**Authors:** Julianne B. Riggs, Eva M. Medina, Loni J. Perrenoud, Diana L. Bonilla, Eric T. Clambey, Linda F. van Dyk, Leslie J. Berg

## Abstract

Investigating the dynamics of virus-host interactions in vivo remains an important challenge, often limited by the ability to directly identify virally-infected cells. Here, we combine detection of a beta-lactamase activated fluorescent substrate with full spectrum flow cytometry to identify primary targets of murine gammaherpesvirus 68 (MHV68) infection in the peritoneal cavity. By optimizing substrate and detection conditions, we were able to achieve multiparameter characterization of infected cells and the ensuing host response. MHV68 infection leads to a pronounced increase in immune cells, with CD8+ T cells increasing by 3 days, and total infiltrate peaking around 8 days post-infection. MHV68 infection results in near elimination of large peritoneal macrophages by 8 days post-infection, and a concordant increase in small peritoneal macrophages and monocytes. Infection is associated with prolonged changes to myeloid cells, with a distinct population of MHC II^high^ large peritoneal macrophages emerging by 14 days. Targets of MHV68 infection could be readily detected. Between 1 to 3 days post-infection, MHV68 infects ~5-10% of peritoneal cells, with >75% being large peritoneal macrophages. By 8 days post-infection, the frequency of MHV68 infection is reduced at least 10-fold, with infection primarily in small peritoneal macrophages, with few infected dendritic cells and B cells. MHV68 infection at 3 days post-infection contains both lytic and latent infection, consistent with the identification of cells with active reporter gene expression. Our findings demonstrate the utility of the beta-lactamase MHV68 reporter system for high throughput single-cell analysis and identify dynamic changes during primary gammaherpesvirus infection.

**Importance:** Identifying virally-infected cells in vivo is key to tracking viral infection and understanding host-pathogen interactions. The ability to further characterize and phenotype virally-infected cells is technically challenging. We use a mouse gammaherpesvirus, MHV68, expressing a reporter gene to identify infected cells during primary infection via flow cytometry. Optimization using this reporter system allowed us to further characterize infected cells via multiparameter full spectrum flow cytometry. Our study provides a technical model for high throughput single-cell immunophenotyping methods in the context of gammaherpesvirus infection. Furthermore, we show that acute MHV68 infection in the peritoneal cavity dramatically changes the immune landscape of this tissue, results in a high number of infected macrophages at early times, and is characterized by both lytic and latent infection within immune cells.

## Introduction

The gammaherpesviruses (γHVs) are a highly conserved family of DNA tumor viruses characterized by their capacity to establish lifelong, latent infection in their hosts. The γHVs include the human viruses, Epstein-Barr virus (EBV) and Kaposi’s-Associated Sarcoma Virus (KSHV), multiple primate γHVs and murine gammaherpesvirus 68 (MHV68, official ICTV nomenclature MuHV-4) (1–3). For most individuals, γHV infection is well-controlled, with no overt deleterious consequences. However, particularly in immunosuppressed individuals, chronic γHV infection is associated with the development of malignancies, including lymphomas and carcinomas (4–6).

The γHVs result in a lifelong infection of their hosts, involving a number of different cell types over time. In vivo, MHV68 is thought to lytically replicate in epithelial cells of mucosal surfaces (e.g. nasopharynx, lung) following mucosal exposure, with long-term latent infection primarily occurring in memory B cells in secondary lymphoid organs such as the spleen (7–9). Additionally, MHV68 has been reported in various tissues depending on the route of infection, [peritoneal cavity (10), lymph node, omentum (11), bone marrow (12), and gut (13–15)], with various types of infection observed in B cell subsets (12, 16–18), myeloid cells (macrophages and dendritic cells) (10, 16), endothelial cells (19), and intestinal epithelial cells (13). The vast array of potential viral reservoirs calls for an efficient and robust method to track infection.

In the past, limiting dilution analyses were used to track MHV68 in various tissues (1). These methods are highly sensitive, but limited in their ability to identify specific cell types that are targets of infection. A variety of MHV68 reporter viruses have been generated. These reporter viruses are able to be used with single-cell immunophenotyping methods, such as flow cytometry and immunohistochemistry. However, these fluorescent proteins are expressed only in the early stages of infection and can be difficult to detect in autofluorescent cell types (20–23).

The use of a beta-lactamase tagged virus, MHV68.LANA:: βlac, expands upon these capabilities (24). The beta-lactamase gene is linked to ORF73, a viral gene encoding LANA, the episome-maintenance protein and transcription factor that is expressed throughout the viral lifecycle. It is notable that while the LANA protein is expressed throughout infection, it is generally expressed at a low level, and thus, combination with the enzymatic amplification of beta-lactamase is critical. Further development of this beta-lactamase tagged virus system allowed tracking of cells infected with panels of mutant viruses both in vitro and in vivo, as well as live cell sorting of rare infected cells for downstream analyses of function and multiparametric mass cytometry and transcriptomics (25, 26).

The peritoneal cavity (PerC) is an important and relevant tissue of immunological function, as well as a source of easily-isolated immune cells. Despite many viral infections being delivered intraperitoneally, the immune response to infection in this cavity is not well characterized. The cavity is composed primarily of macrophages and B cells (27). Peritoneal B cells are predominantly B-1 cells, “innate” B cells that produce natural antibodies (28). MHV68 has been identified to establish latency in peritoneal macrophages and B-1 cells (10, 17). Macrophages are of two types: large peritoneal macrophages (LPMs), characterized as tissue resident, self-renewing “clean-up” phagocytes, and small peritoneal macrophages (SPMs), which are more inflammatory, and replaced from circulating monocytes in the blood (29). LPMs are the dominant population, however it is SPMs that are elicited from thioglycolate stimulation, a common method in obtaining peritoneal macrophages (29). Lytic infection of primary macrophages has also been established *in vitro* (21, 30) but characterization of acute gammaherpesvirus infection of the PerC is limited.

Here, we optimized detection of a reporter virus, MHV68.LANA::βlac, using full spectrum (i.e. spectral) flow cytometry to achieve single cell resolution of virus infection during primary infection in the peritoneal cavity. We show that MHV68 infection in the peritoneal cavity leads to dramatic changes in immune cell populations, most notably a transient elimination of large peritoneal macrophages and an increase in small peritoneal macrophages, monocytes, and CD8+ T cells. We use MHV68.LANA::βlac to track infection over time, identifying and immunophenotyping virally infected cells up to two weeks post infection. We show that large peritoneal macrophages represent the dominant reservoir of MHV68 during acute infection, and that infection in the peritoneal cavity at this time is a mixture of both lytic and latent infection. These studies demonstrate a method for expansive immunophenotyping that can be integrated with detection of the MHV68.LANA::βlac reporter virus, and present new insights in acute gammaherpesvirus infection.

## Methods

### Mice

C57BL/6 mice were purchased from Jax and bred in house (Catalog #00664). Mice were maintained at the University of Colorado Anschutz Medical Campus in accordance with IACUC protocol and were used for experiments at 6-12 weeks of age. Both male and female mice were used.

### Viral infections

Wild-type (WT) MHV68 and MHV68.LANA::βlac viruses were grown and prepared as described previously (25). For *in vitro* infections, mouse 3T12 fibroblasts and *ex vivo* peritoneal cells were infected overnight, with no removal of inoculum, at a multiplicity of infection (MOI) of 1 and 10 plaque forming units (PFU) per cell, respectively. Cells were cultured in DMEM with L-glutamine, 1% Penicillin/Streptomycin and 10% FBS. For *in vivo* infections, mice were inoculated intraperitoneally (i.p.) with 1×10^6^ PFU of virus diluted in sterile PBS.

### Harvest of peritoneal cells

Mice were sacrificed via CO_2_ euthanasia and cervical dislocation. The peritoneal cavity was exposed and filled with 10mL of ice cold PBS containing 3% FBS using a 27 gauge needle. After removal of the needle, the mouse was then agitated to encourage release of immune cells from the peritoneal wall, followed by fluid and cell extraction using an 18 gauge needle. If necessary, cells were subjected to red blood cell lysis with ACK lysis buffer (Gibco Catalog #A1049201), washed and counted using the Beckman Coulter Vi-CELL BLU.

### Flow cytometric analysis of peritoneal cells

Following harvest and preparation, peritoneal cells were subjected to flow cytometric analysis. First, cells were assessed for viability using a 30 min RT incubation with Zombie NIR (BioLegend) diluted in PBS. CCF2-AM substrate (LiveBLAzer™ FRET-B/G Loading Kit with CCF2-AM, ThermoFisher) (31) was freshly diluted prior to substrate loading for 30-60 minutes RT in PBS. Cells were subjected to Fc receptor blocking using Fc Shield (Tonbo Biosciences) for 10 minutes, followed by antibody staining for 25 minutes. The following antibodies, with antibody clones designated in parenthesis, were used: CD19(1D3)-PECy7, CD11b(M1/70)-APC (eBiosciences/ThermoFisher), CD11c(HL3)-BUV737, CD4(GK1.5)-BUV395, CD8a(53-6.7)-PerCPCy5.5 (BD), CD3e(145-2C11)-PECy5, F4/80(BM8)-BV785, MHC II(M5/14.15.2) PerCP, Gr1(RB6-8C5)-AlexaFluor700, Gal3(M3/38)-AlexaFluor647 (BioLegend). All staining reactions were done at RT in the dark in PBS for antibody stains and PBS containing 1% FBS for Fc receptor blockade. Samples were acquired on a Cytek Aurora Spectral Analyzer with five lasers (UV, violet, blue, yellow-green, and red) using SpectroFlo software.

### MHV68 Reactivation Assay

C57BL/6 mouse embryonic fibroblasts (MEFs) were cultured in DMEM with L-glutamine, 20% FBS, 1% Penicillin/Streptomycin and 1x Amphotericin B at 5,000 cells per well in 96-well flat bottom tissue culture-treated plates. Peritoneal cells (PerCs) were harvested from mice inoculated with PBS alone, MHV68 or MHV68.LANA::βlac at 3 days post infection (dpi).. PerCs were washed, counted, and diluted to the indicated concentrations. In parallel, cells were subjected to mechanical disruption and plated in a comparable series of dilutions prior to plating on MEF monolayers (32). Co-cultures were incubated at 37 C for 21 days prior to assessment of cytopathic effect (CPE).

### Data Analysis

Analysis of flow cytometry data was performed using Cytek SpectroFlo version 2.1 and DeNovo FCS Express version 7. Fluorophore signatures with partial overlap of emission spectra were resolved by unmixing using SpectroFlo software without compensation. Statistical analysis was performed using student’s t test in GraphPad Prism version 8.

## Results

### Optimization of MHV68.LANA::βlac detection analyzing CCF2-AM substrate cleavage by spectral flow cytometry

We sought to harness this powerful technology for multiparametric flow cytometry with the beta lactamase reporter virus to track virally infected cells *in vitro and in vivo.* Previous use of a beta-lactamase expressing MHV68 virus detected virally-infected cells utilizing conventional, band-pass flow cytometry (12, 24, 25, 32). Though these studies were informative, this technique collects narrow windows of fluorescence emission. Given the broad emission spectrum conferred by the CCF2-AM fluorescent substrate, previous studies were limited in the number of fluorophores that could be simultaneously measured. Recently, full spectrum (herein referred to as spectral flow cytometry) has been developed, a technique that characterizes fluorophores by their entire spectra, rather than peak emission, using more sensitive light detectors. Analysis of whole emission spectra allows for the combination of fluorophores relatively close in peak emission, vastly expanding the number of fluorophores that can be simultaneously analyzed. Full spectrum flow cytometry also allows for the detection of fluorescent reporters and proteins whose spectra are not well characterized or are unknown (33, 34). To date, full spectrum flow cytometry has primarily been applied to conventional staining reagents to significantly increase the number of parameters for simultaneous analysis (e.g. (35).

Studies with the MHV68.LANA::βlac reporter virus rely on detection of cleavage of a beta-lactamase substrate, such as CCF2-AM (31, 36), an esterified beta-lactamase substrate that readily crosses the cell membrane (Figure 1). Once inside the cell, endogenous cytoplasmic esterases cleave ester groups, resulting in the lipophobic molecule CCF2. CCF2 consists of a coumarin (identified as “A” in Figure 1A) linked to a fluorescein (identified as “B”, Figure 1A) via a cephalosporin group. The coumarin group absorbs light at a maximum wavelength of 408 nm and, via Fluorescence Resonance Energy Transfer (FRET), transfers the energy to the fluorescein group, which emits at a maximum wavelength of 520 nm. In the uncleaved state, CCF2 (Figure 1A, left) exhibits peak emission from the fluorescein group. In the presence of beta-lactamase enzymatic activity (Figure 1A, right), the cephalosporin group is cleaved to separate the coumarin and fluorescein groups, disrupting FRET and altering the resulting fluorescence to the coumarin group, which emits at a maximum wavelength of 447 nm. Thus, the presence of beta-lactamase (βlac) enzyme, such as the LANA::βlac fusion protein, can be detected by the unique fluorescence profile of the cleaved CCF2 substrate.

**Figure 1:**
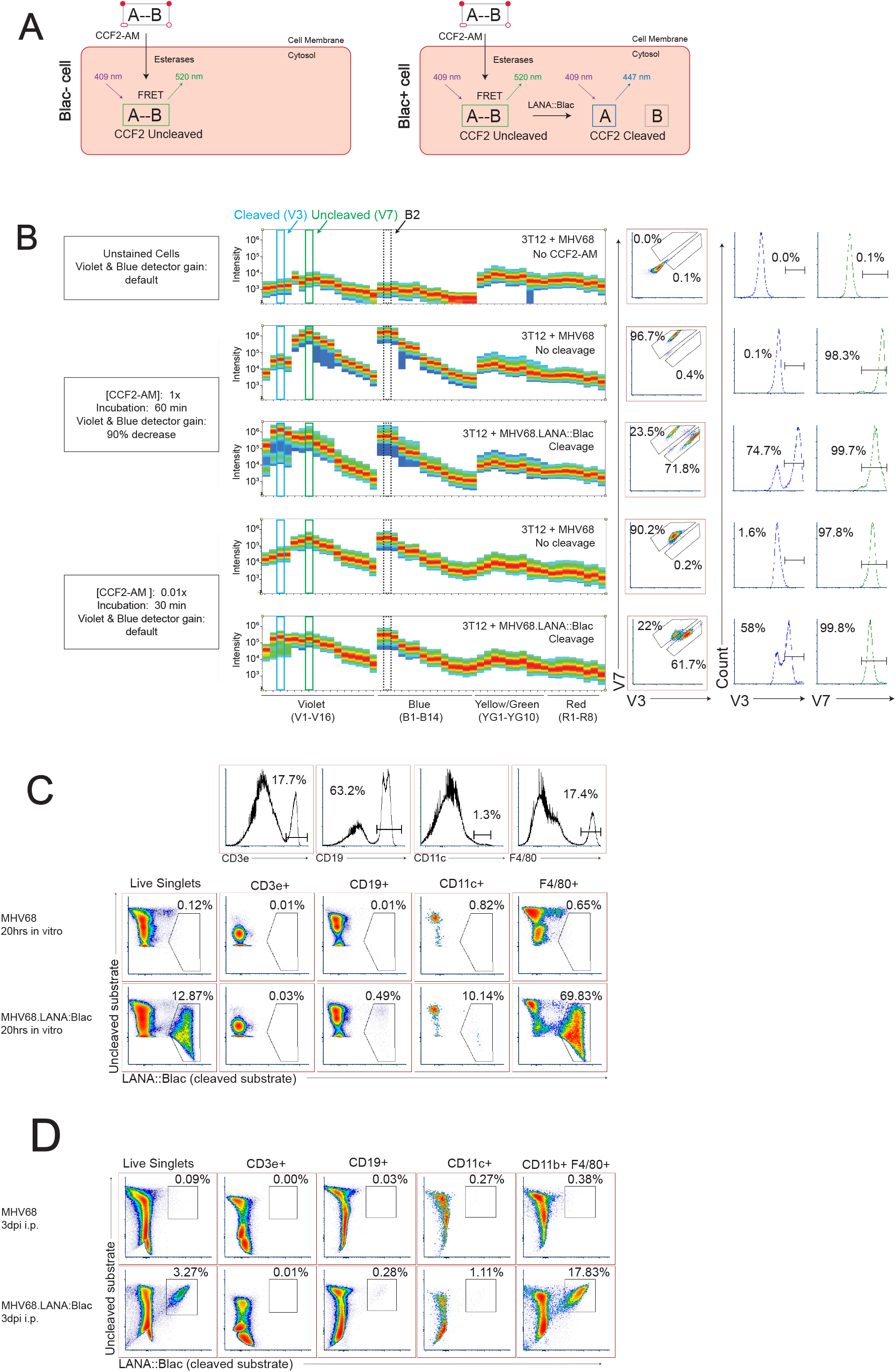
Optimization of MHV68.LANA::βlac detection and CCF2-AM cleavage by full spectrum flow cytometry. (A) Graphic depicting CCF2-AM entrance, processing and fluorescence in cells in the absence (left) or presence (right) of mLANA::βlac. “A” represents the coumarin component of CCF2, while “B” represents fluorescein. (B) CCF2-AM titration and detection in 3T12 mouse fibroblasts, infected with either MHV68 (i.e. no CCF2-AM cleavage) or MHV68.LANA::βlac (i.e. CCF2-AM cleavage) and incubated with the indicated concentration of CCF2-AM at 20 hours post-infection. Shown are full emission spectra from stained cells. Each row depicts fluorescence intensity under different experimental conditions (MHV68 or MHV68.LANA::βlac infection), varying CCF2-AM concentration and incubation time, and gain for Violet and Blue detectors of a Cytek Aurora spectral analyzer. Top row: unstained fibroblasts. Middle rows: fibroblasts subjected to manufacturer’s recommended conditions for CCF2-AM staining. Bottom rows: fibroblasts subjected to optimized, titrated CCF2-AM staining. Each row depicts from left to right: fluorescent spectral intensity across 48 detectors, with fluorescence emission in the V3 and V7 detectors highlighted by dot plot and histogram. CCF2-AM associated fluorescence for the uncleaved substrate is detected in V7, with cleaved substrate detected in V3. (C) Dot plots showing optimized staining conditions in infected mouse peritoneal cells (PerCs) in vitro. PerCs were extracted and infected overnight in vitro with MHV68 or MHV68.LANA::βlac. 20 hrs later, cells were stained with CCF2-AM as well for viability and surface markers. Histograms are shown to demonstrate how populations were defined. (D) Dot plots showing optimized staining conditions on infected PerCs in vivo. Mice were infected i.p. with 1 million PFUs of MHV68 or MHV68.LANA::βlac. Three days after infection, mice were sacrificed and PerCs were harvested and stained with CCF2-AM as well as for viability and surface markers. Populations shown were gated on live singlets. Data representative of three experiments, with 2-3 mice per condition per experiment.

Despite the established utility of the MHV68.LANA::βlac reporter system (24, 25), the broad emission spectrum of the cleaved CCF2 substrate has prevented robust multiparameter flow cytometric analysis on a conventional flow cytometer (data not shown). With the recent establishment of full spectrum, spectral flow cytometry, we sought to test whether it may be possible to further empower flow cytometric analysis of the βlac reporter system. To do this, we infected mouse 3T12 fibroblasts with either WT MHV68, which lacks βlac expression, or MHV68.LANA::βlac for 20hrs at an MOI of 1 PFU/cell, and incubated cells with a final concentration of 10 μM CCF2-AM substrate according to manufacturer’s recommendations. When cells were analyzed for fluorescence on a Cytek Aurora spectral analyzer, both MHV68 and MHV68.LANA::βlac infected samples were characterized by an extremely bright fluorescent signal (Figure 1B), consistent with what had been observed in previous reports with conventional flow cytometry (ETC, personal communication). Maximal fluorescence of the uncleaved CCF2 substrate, in MHV68-infected cultures, was detected in the V7 and B2 fluorescent detectors (Figure 1B, second row), consistent with fluorescein emission. In contrast, MHV68.LANA::βlac-infected cultures demonstrated an additional fluorescence maximum in the V3 fluorescent detector, consistent with LANA::βlac-dependent substrate cleavage. We propose that retention of fluorescence in the V7 and B2 detectors likely results from incomplete cleavage of CCF2 in LANA::βlac+ cells, a phenomenon also observed in other reports using this system (24, 25, 31). Despite the ability to detect a unique fluorescent signature in MHV68.LANA::βlac-infected cells, CCF2-associated fluorescence in these conditions was so great that the original Cytek assay gain settings for the violet and blue detectors had to be decreased by more than 90%. This reduction severely compromised the potential to analyze additional fluorophores excited primarily by these lasers.

To address this limitation, we analyzed the impact of CCF2-AM substrate concentration and incubation time on fluorescent emission. Notably, use of 0.1 μM CCF2-AM, a 100-fold dilution in substrate, produced a signal that was readily detectable and on-scale yet did not require altering laser voltages on the cytometer (Figure 1B, bottom two rows). We further determined that incubation with the CCF2-AM substrate for half the recommended time (30 minutes instead of 60 minutes) still produced a robust signal (Figure 1B). These optimized substrate conditions afforded the opportunity to discriminate between uncleaved substrate, resulting in maximal fluorescence in the V7 and B2 detectors, and cleaved CCF2 substrate, uniquely fluorescing in the V3 detector while retaining fluorescence in the V7 and B2 detectors. Significantly, these settings did not require compromised gains for either the blue or violet laser, allowing detection of other fluorophores. One important aspect of spectral flow cytometry is the accurate definition of the unique emission spectrum for each fluorophore. In order to accurately unmix the cleaved CCF2 fluorescent signature from MHV68.LANA::βlac infected samples, we used an iterative process in which cells with cleaved CCF2 fluorescence from MHV68.LANA::βlac infected samples were identified, exported and reimported to define an accurate cleaved CCF2 spectrum (details outlined in Supplementary Figure 1).

To determine if our optimized CCF2-AM staining conditions worked in primary cells, we infected mouse peritoneal cells (PerCs) directly *ex vivo*. The peritoneal cavity is rich in macrophages and B cells, two cell types known to be latently infected following MHV68 infection (10, 17). PerCs were isolated and incubated overnight *in vitro* with either WT MHV68 or MHV68.LANA::βlac (MOI = 10 PFUs/cell), followed by use of optimized CCF2-AM incubation conditions. Wild-type (WT) MHV68 infected cultures had minimal fluorescence in the CCF2 cleaved substrate detection channel (Figure 1C, top row). In contrast, ~12% of MHV68.LANA::βlac infected PerCs demonstrated CCF2 cleaved substrate-dependent fluorescence (Figure 1C, bottom row). Furthermore, these conditions allowed us to use additional fluorophores excited by the violet and blue lasers, a result unachievable using the manufacturer’s recommended staining conditions. The use of multiple antibodies to detect a panel of cell surface markers allowed us greater resolution to immunophenotype virus-positive cells. MHV68.LANA::βlac was detected in a high frequency of F4/80+ macrophages, with a lower frequency of cleaved substrate detected in CD11c+ dendritic cells and CD19+ B cells, and no detectable fluorescence in CD3+ T cells (Figure 1C). Next, we tested our ability to detect MHV68.LANA::βlac infection in PerCs harvested from mice that were subjected to intraperitoneal infection and harvested at 3 days post-infection (dpi). Again, a robust cleaved CCF2 signal was observed, particularly among F4/80+ macrophages (Figure 1D).

For the above experiments, samples were collected on the same day as CCF2-AM staining. We also found that samples stained with CCF2-AM could be subjected to overnight fixation, without any degradation of CCF2-dependent fluorescence even after 20 hrs (Supplementary Figure 2). Notably, however, samples stained with CCF2-AM were not tolerant to permeabilization, as this caused pronounced loss of signal for both uncleaved and cleaved CCF2 (Supplementary Figure 2). With optimization established, we sought to apply this to analyze primary MHV68 infection using MHV68.LANA::βlac infection and spectral flow cytometry.

### MHV68 results in dramatic changes to the immune landscape of the peritoneal cavity

The viral lifecycle of γHVs is dynamic, with significant changes in viral load and cell and tissue tropism over time. To identify the consequence of acute MHV68 infection on cell types and targets of virus infection, mice were infected intraperitoneally (i.p.) with either WT MHV68 or MHV68.LANA::βlac, and analyzed over 14 days of infection.

Cell counts revealed that i.p. MHV68 infection triggers a robust inflammatory response in the peritoneal cavity, peaking at 8 dpi (Figure 2A). This is accompanied by appreciable splenomegaly (Supplementary Figure 3). Leukocyte cell counts began to subside after 8 dpi, but remain elevated compared to mock-infected mice. MHV68 infection induced a prominent increase in T cell frequency and number, dominated by CD8+ T cells (Figures 2B and 2C). Notably, CD8 T cells increased by 3dpi, indicating rapid engagement of CD8 T cells. Immune cell populations were defined as according to the gating strategy detailed in Supplementary Figure 4 (Supplementary Figure 4).

**Figure 2:**
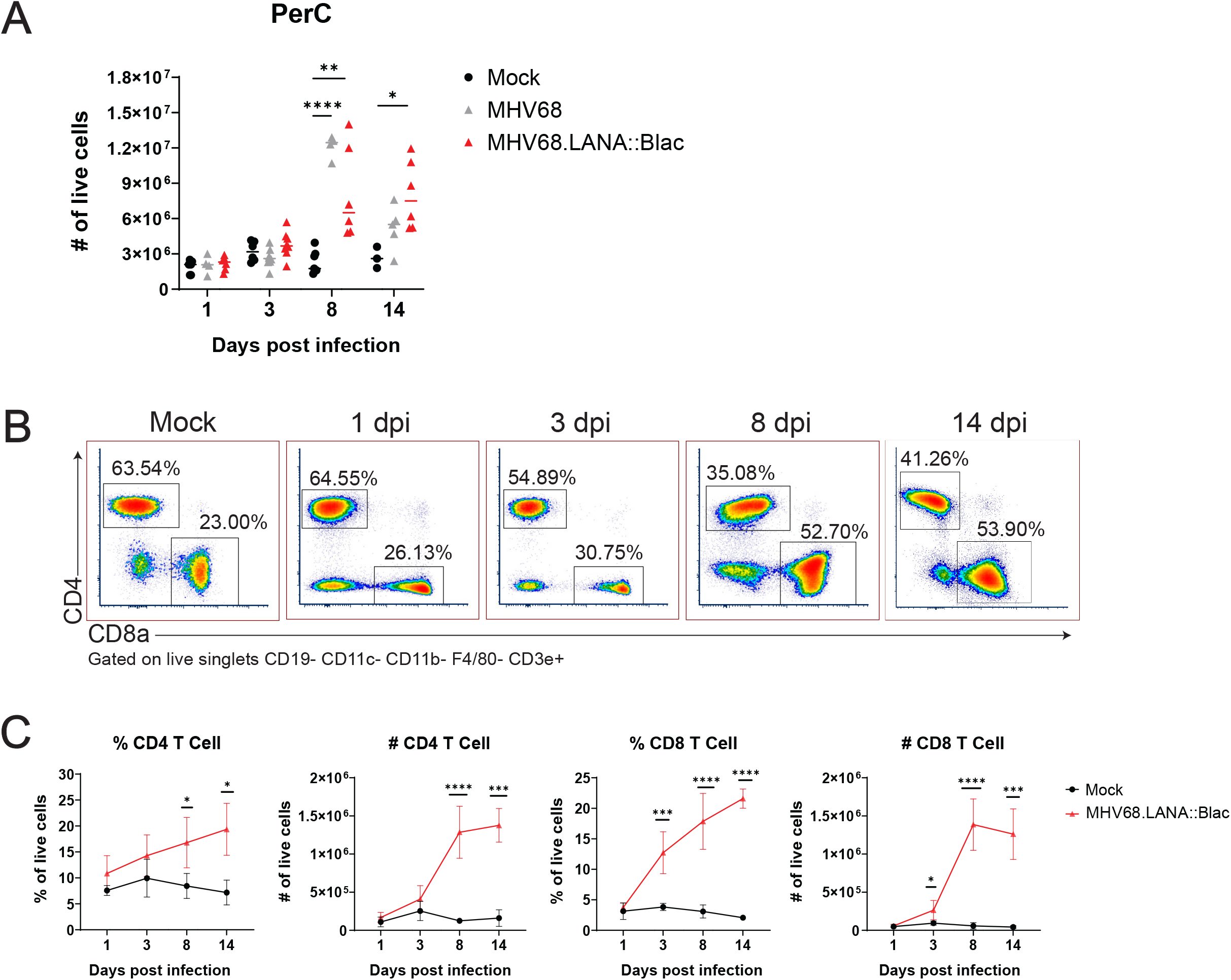
Intraperitoneal MHV68 infection induces T cell accumulation in the peritoneal cavity. Experimental conditions were performed as outlined in Figure 1D. Mice were harvested at 1, 3, 8 and 14 dpi. (A) Viable cell counts of PerCs as a function of time and infection cohort. Each symbol is representative of one mouse and bars represent the mean value of samples. (B) Flow cytometry dot plots showing percent of CD4- and CD8a-positive PECs at various time points post infection. Samples are gated on live, singlets CD19- CD11c- CD11b- F4/80- CD3e+ cells. (C) Quantification of flow cytometry data showing mean percent and number of CD4 and CD8a-positive PECs are various time points post infection. Statistical analysis was performed using student’s t test in GraphPad Prism (*p<0.05, **p<0.01, ***p<0.001,****p<0.0001). Data representative of three experiments with 2-3 mice per condition per experiment.

MHV68 infection was also associated with pronounced changes in myeloid subsets. In naïve mice, there are two dominant macrophage populations, large peritoneal macrophages (LPM), defined as F4/80^High^ CD11b^High^ MHC II^Low^, and small peritoneal macrophages (SPM), defined as F4/80^Mid^ CD11b^Mid^ MHC II^High^ (29). MHV68 infection reduced the frequency of LPMs relative to mock-infected mice, from day 1 through day 14 post-infection, with no LPMs detected at 8 dpi and an apparent reappearance by day 14 (Figure 3A-B). This dramatic loss of LPMs was accompanied by a pronounced increase in the frequency and number of F4/80^Mid^ CD11b^Mid^ cells, consistent with an SPM or monocyte phenotype, between days 3 to 14 post-infection (Figure 3A-B). The phenotype of F4/80^Mid^ CD11b^Mid^ cells changed over time. In mock-infected mice, F4/80^Mid^ CD11b^Mid^ cells were predominantly a Gr1^Low^ MHC II^High^ phenotype consistent with SPMs (29). Following MHV68 infection, however, F4/80^Mid^ CD11b^Mid^ cells demonstrated a transient increase in the frequency of Gr1^High^ MHC II^Low^ cells, consistent with monocyte infiltration into the peritoneal cavity (Figure 3C). By day 8 and 14 post-infection, F4/80^Mid^ CD11b^Mid^ cells were primarily Gr1^Low^ MHC II^High^ cells, consistent with SPMs (Figure 3C).

**Figure 3:**
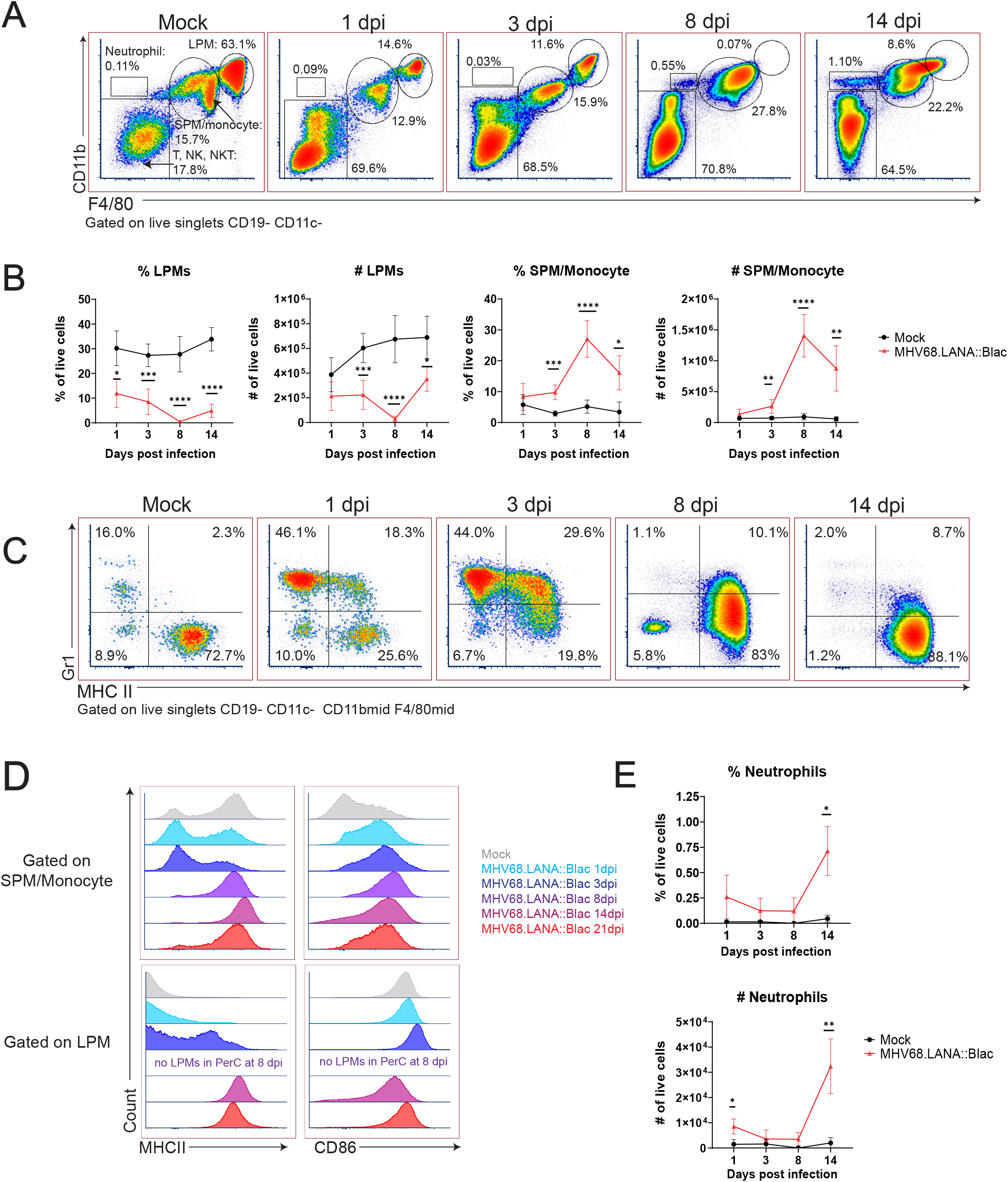
Alterations in peritoneal myeloid subsets after MHV68 infection. Flow cytometric analysis of myeloid cells and neutrophils in the peritoneal cavity after MHV68 infection. (A) Flow cytometric analysis of PerCs, showing expression of CD11b and F4/80 on live, singlet, CD19- CD11c- events at the indicated times post infection. Gates identify distinct cell subsets, including large peritoneal macrophage (LPM) and small peritoneal macrophages (SPM). (B) Quantification of flow cytometry data shown in (A) showing mean percent and number of live singlets that are LPMs and SPM/monocytes at various times post infection. (C) Dot plots showing expression of Gr-1 and MHC II on CD19- CD11c- CD11b^mid^ F4/80^mid^ PerCs over time. (D) Histograms comparing expression of MHC II (left) and CD86 (right) in SPMs/monocytes (top) and LPMs (bottom), as a function of time (colored according to legend). Day 8 samples did not have any detectable LPMs (as indicated). (E) Quantification of flow cytometry data shown in (A), demonstrating mean percent and number of neutrophils at time points post infection. Statistical analysis was performed using student’s t test in GraphPad Prism (*p<0.05, **p<0.01, ***p<0.001,****p<0.0001). Data representative of three experiments with 2-3 mice per condition per experiment.

The disappearance and reappearance of the LPM population by 14 dpi suggested self-renewal and/or replenishment. Despite prior reports detailing LPMs as self-renewing (37), the complete absence of LPMs at 8 dpi raised the possibility that SPMs and/or monocytes from the circulation enter the peritoneal cavity and acquire the characteristics of LPMs. Notably, as infection progressed, SPM/monocyte and LPM populations became less distinct based on CD11b and F4/80 expression (e.g. see 14 dpi, Figure 3A). SPM and LPM protein expression was further altered by infection. In uninfected mice, SPMs have been characterized as MHC II^High^ CD86^Low^, in contrast to LPMs which are MHC II^Low^ CD86^High^ (29); we also find these distinct phenotypes in mock-infected mice (Figure 3D). In contrast, by 14 and 21 dpi, F4/80^Mid^ CD11b^Mid^ SPM/monocytes upregulated CD86 expression compared to mock-infected conditions (Figure 3D). Conversely, F4/80^High^ CD11b^High^ LPMs increased their expression of MHC II, with LPMs demonstrating an MHC II^High^ phenotype by day 14 and 21 dpi (Figure 3D). These phenotypic changes resulted in SPM/monocyte and LPM populations both expressing an MHC II^High^ CD86^High^ phenotype by 14 dpi, blurring the phenotypic distinction between these cell subsets. In addition to phenotypic changes in macrophage subsets in the peritoneal cavity, MHV68 infected mice further demonstrated an increase in the frequency of neutrophils in the peritoneal cavity by 14 dpi (Figure 3E). These data demonstrate multiple time-dependent changes in leukocyte composition and phenotype following primary MHV68 infection.

### MHV68.LANA::βlac+ is detectable in macrophages, dendritic cells and B cells in the peritoneal cavity

We next analyzed the cellular distribution of MHV68.LANA::βlac infection in the peritoneal cavity over time. The frequency of LANA::βlac+ cells was relatively constant between 1 dpi and 3 dpi, with at least 5% of total peritoneal cells demonstrating CCF2-AM substrate cleavage, a single-cell marker for LANA::βlac expression (Figure 4). The frequency of LANA::βlac+ cells decreased significantly (~10-fold) by 8 dpi, with a further decrease in the frequency of LANA::βlac+ PerCs by 14 dpi (Figure 4). Despite the decreased frequency of LANA::βlac+ cells over time, these cells were readily detected above the limit of detection defined by coincident analysis of mock and WT MHV68 infected samples (Figure 4).

**Figure 4:**
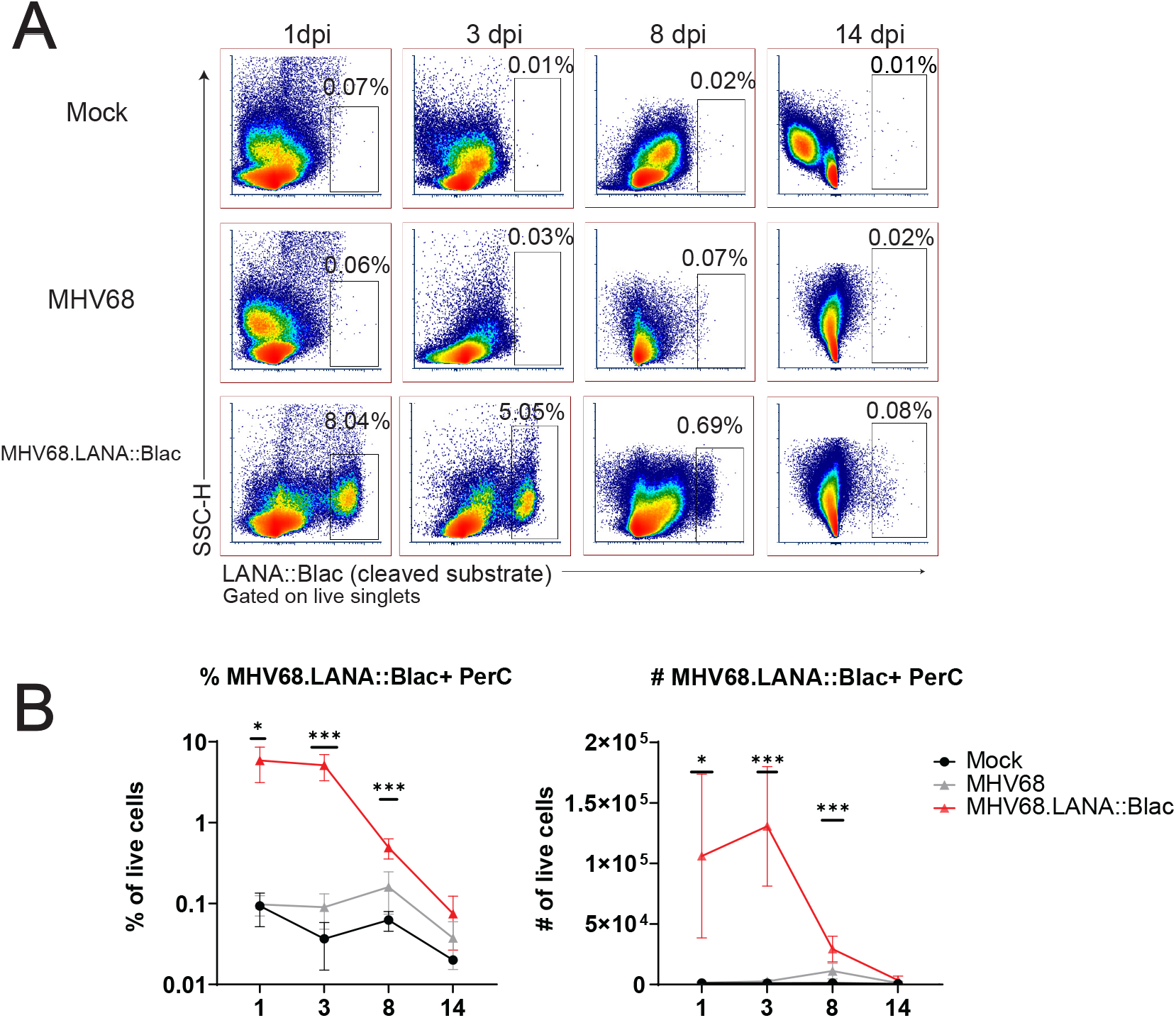
MHV68.LANA::βlac is detectable during acute infection in the peritoneal cavity. Flow cytometric analysis of MHV68.LANA::βlac infection in the peritoneal cavity of MHV68 infected mice. (A) Dot plots showing the frequency of PerCs positive for cleaved CCF2 fluorescence throughout infection, comparing peritoneal cells isolated from mock, MHV68 or MHV68.LANA::βlac infected mice. Populations were gated on live singlets. (B) Quantification of the frequency and number of LANA::βlac+ PerCs defined based on positive cleaved CCF2 fluorescence. Statistical analysis was performed using student’s t test in GraphPad Prism (*p<0.05, **p<0.01, ***p<0.001,****p<0.0001). Data representative of three experiments with 2-3 mice per condition per experiment.

We next immunophenotyped PerCs from infected mice using a panel of 12 cell surface markers. As predicted from our initial *in vivo* experiments (Figure 1D), we detected the virus in multiple cell types, including macrophages, dendritic cells, and B cells, at varying frequencies (Figure 5). Of these, the majority (>75%) of LANA::βlac+ cells were identified within large peritoneal macrophages early after infection (1 dpi and 3 dpi) (Figure 5). By 8 dpi, however, F4/80^Mid^ CD11b^Mid^ small peritoneal macrophages were the major LANA::βlac+ population (Figure 5), a change in tropism coincident with a nearly complete disappearance of LPMs from the peritoneal cavity (as shown in Figure 3). B cells and dendritic cells accounted for less than 5% of LANA::βlac+ cells through 14 dpi (Figure 5). Thus, we conclude that macrophages are the major target of MHV68 infection in the peritoneal cavity during the first two weeks of infection.

**Figure 5:**
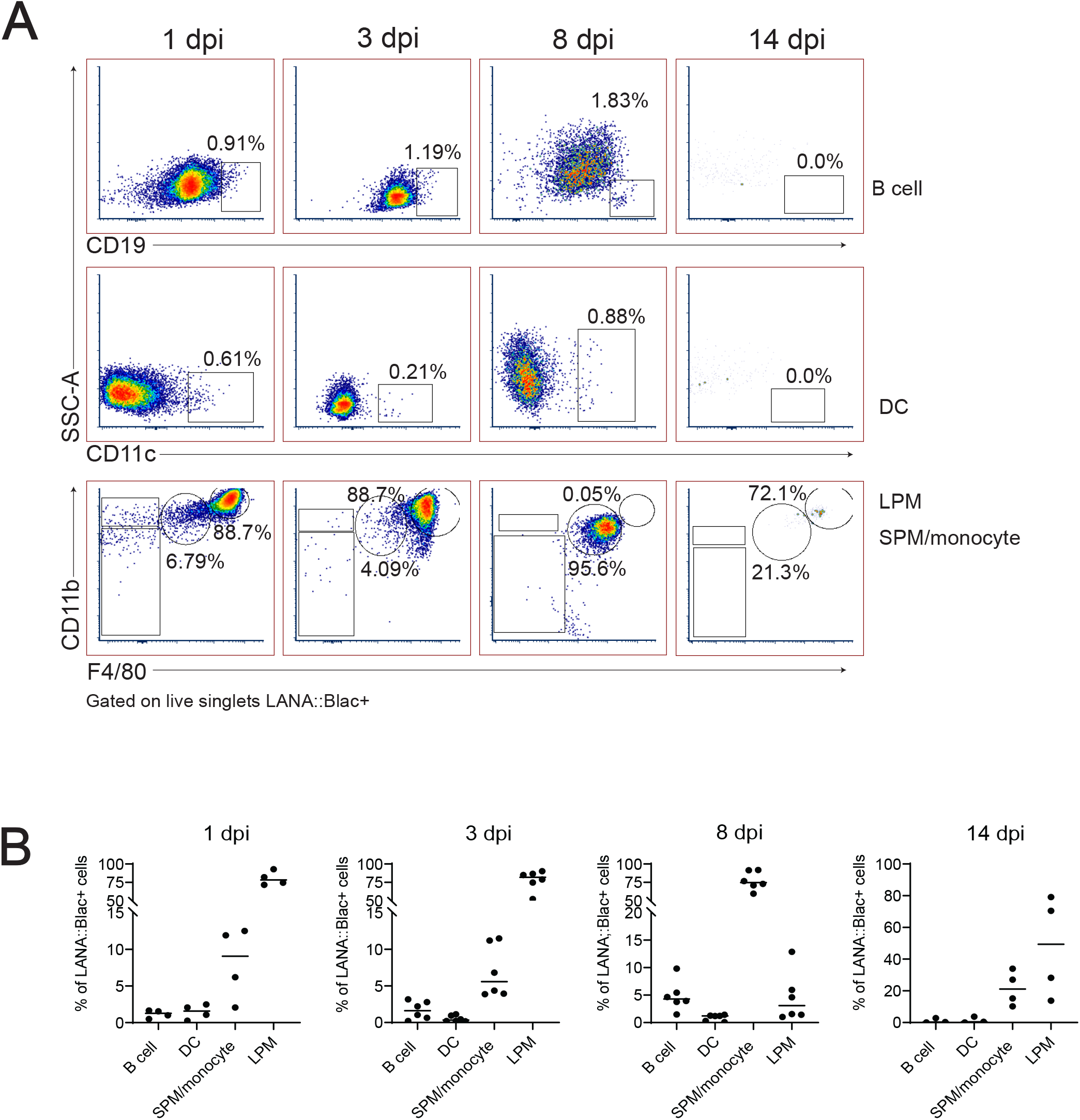
Predominant detection of MHV68.LANA::βlac infection in macrophages in the peritoneal cavity. Flow cytometric analysis of MHV68.LANA::βlac infection within different leukocyte subsets in the peritoneal cavity of MHV68.LANA::βlac infected mice. (A) Dot plots quantifying the frequency of LANA::βlac+ events (as defined in Figure 4) that are CD19+ B cells (top row), CD11c+ dendritic cells (middle row), or CD11b^High^ F4/80^High^ LPMs or CD11b^Mid^, F4/80^Mid^ SPM/Monocytes (bottom row). (B) Quantitation of the frequency of LANA::βlac+ (i.e. cleaved CCF2+ cells) that are B cells, DCs, SPM/monocytes, or LPMs at various times post infection. Each dot is representative of one mouse, and lines represent mean of samples. Data representative of three experiments with 2-3 mice per condition per experiment.

In parallel, we assessed the percentage of each cell population that was virally-infected. This analysis identified a hierarchy of infection 1 day post infection, with ~25% of LPMs, 7% of dendritic cells, 5% of SPM/monocytes and 0.2% of B cells which were LANA::βlac+ (Figure 6). LPMs remained a predominant target of infection at 3 dpi, with >40% of LPMs characterized by LANA::βlac expression, in contrast to SPM/monocytes, dendritic cells and B cells which contained <3% LANA::βlac+ cells out to 14 dpi (Figure 6). LANA::βlac expression was not detected in T cells at any time analyzed. These data identify multiple targets of primary MHV68 infection in the peritoneal cavity.

**Figure 6:**
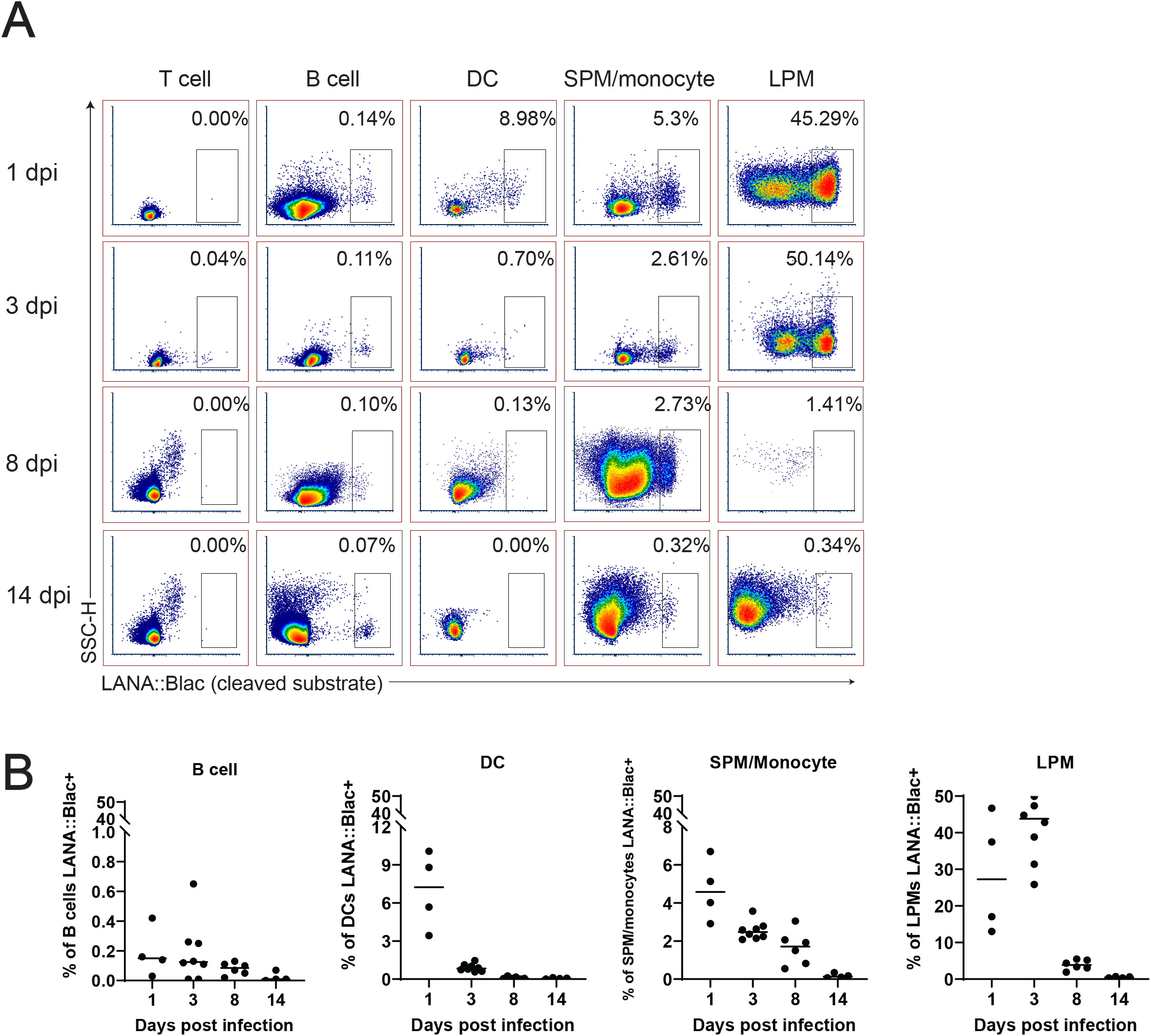
A substantial portion of peritoneal macrophages are positive for MHV68.LANA::βlac infection during acute infection. Flow cytometric analysis of cell subsets that demonstrate MHV68.LANA::βlac infection in the peritoneal cavity of MHV68.LANA::βlac infected mice. (A) Dot plots showing the frequency of T cells, B cells, dendritic cells, SPM/monocytes and LPM that express cleaved CCF2 fluorescence, the indicator for LANA::βlac expression, at various times post infection. (B) Graphs demonstrating data shown in (A). Each dot is representative of one mouse and lines represent mean of samples. Data representative of three experiments with 2-3 mice per condition per experiment.

### MHV68 infection is associated with altered macrophage expression of MHC II and CD86 in infected and uninfected cells

Given the impact of MHV68 infection on myeloid cells, and the partial infection of LPMs and SPMs, we next analyzed how MHV68 infection affected macrophage phenotype in a cell-intrinsic and cell-extrinsic manner. To do this, we analyzed MHC II and CD86 expression, two proteins differentially expressed in LPMs and SPMs at baseline (29), comparing LPMs and SPMs in mock-infected mice with LANA::βlac+ (virus-infected) and LANA::βlac- (uninfected) cell subsets from MHV68 infected mice. As anticipated, mock-infected mice contained LPMs with a predominant MHC II^Low^ CD86^High^ phenotype (Figure 7, “Mock LPM”). After 1 dpi, >94% of LANA::βlac+ and LANA::βlac-LPMs were MHC II^Low^ CD86^High^, comparable to mock-infected LPMs (Figure 7, left panel). By 3 dpi, however, MHV68 infected mice demonstrated an increased frequency of MHC II^High^ LPMs compared to mock-infected mice. MHC II^High^ CD86^High^ LPMs were most prominent among virally-infected (LANA::βlac+) LPMs (49.2% of events), with an intermediate frequency (23.6%) in LANA::βlac-LPMs (Figure 7, middle panel). CD86 expression remained high in LPMs in all conditions (Figure 7, bottom panel). These data demonstrate that MHV68 infection is associated with an increased frequency of MHC II^High^ LPMs, a process that occurs in both virus-infected and ‒uninfected cells during acute infection.

**Figure 7:**
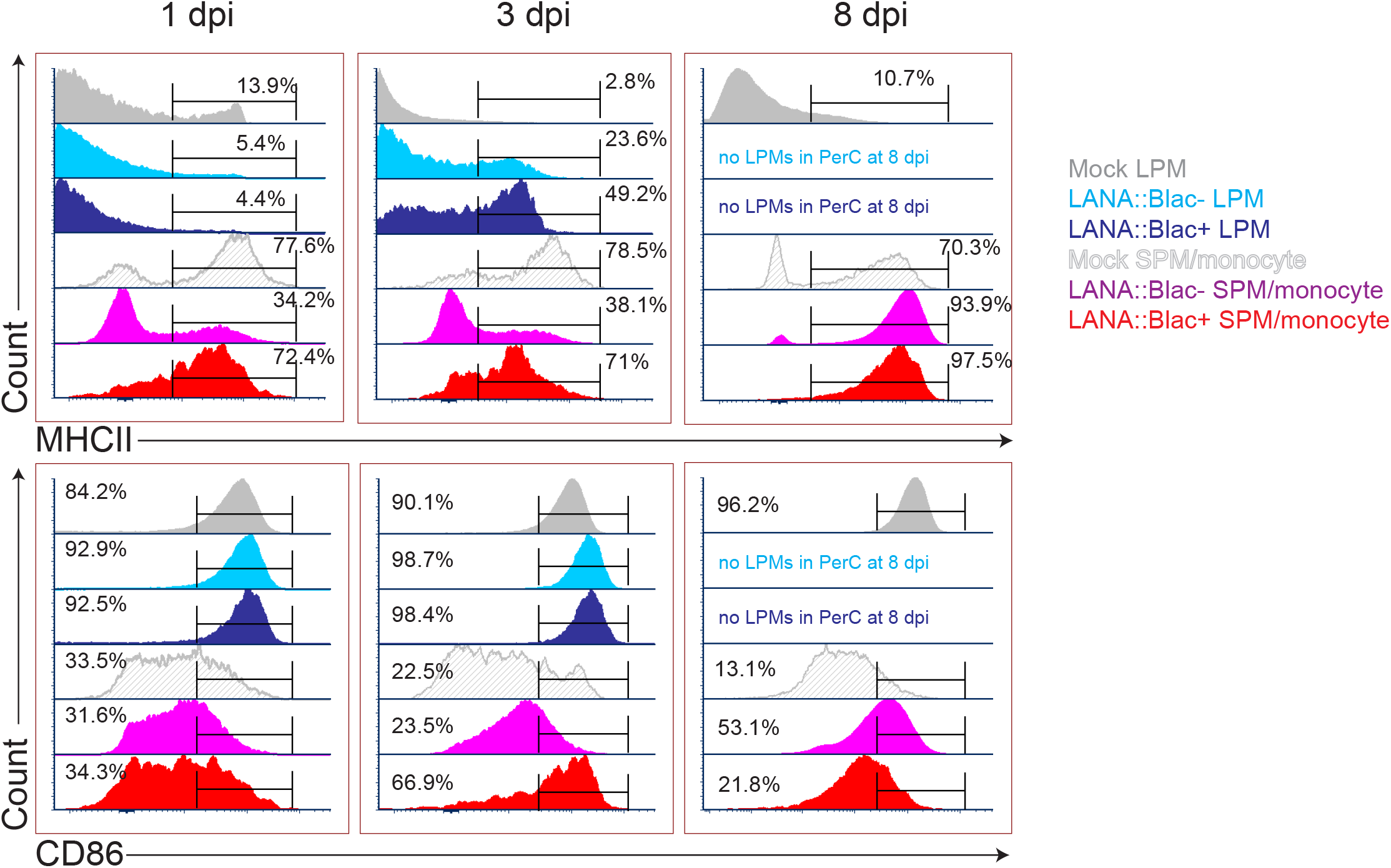
MHV68 infection is associated with altered macrophage expression of MHC II and CD86 in infected and uninfected cells. Flow cytometric analysis of MHC II (top row) and CD86 (bottom row) expression in large (LPM) and small (SPM) peritoneal macrophages, in mock or MHV68.LANA::βlac infected PerCs. Histograms compare the phenotype of virus-infected (LANA::βlac+) and uninfected (LANA::βlac-) cells. Gates define the percent of events positive for each marker. Virus-infected (LANA::βlac+) and uninfected (LANA::βlac-) cells were defined according to the gates drawn in Figure 6A. Data representative of three experiments with 2-3 mice per condition per experiment.

In parallel, we analyzed the impact of MHV68 infection on the phenotype of F4/80^Mid^ CD11b^Mid^ cells, a phenotype containing a mixed population of SPM and monocytes (referred to collectively as SPM/monocytes). Mock-infected SPM/monocytes were primarily characterized by an MHC II^High^ CD86^Low^ phenotype (Figure 7). After overnight infection and through 3 dpi, LANA::βlac+ SPM/monocytes had a relatively comparable frequency of MHC II^High^ events comparable to mock-infected mice. In contrast, LANA::βlac-SPM/monocytes at these early times had a reduced frequency of MHC II^High^ events compared to mock-infected cells (Figure 7). By 8 dpi, however, both LANA::βlac+ and LANA::βlac-SPM/monocytes had an increased frequency of MHC II^High^ events relative to mock-infected mice. While CD86 expression was comparable in all measured SPM/monocyte populations after overnight infection, LANA::βlac+ demonstrated a transient upregulation at 3 dpi relative to mock-infected and LANA::βlac-cells, with both LANA::βlac+ and LANA::βlac-cells characterized by a modest upregulation of CD86 by 8 dpi (Figure 7). In total, these data identify that MHV68 infection is associated with enhanced MHC II and CD86 expression in virally-infected and ‒uninfected macrophage subsets in the peritoneal cavity.

### The MHV68-infected peritoneal cavity contains both lytic and latent infection early after infection

Our analysis of the cellular distribution of LANA:: βlac identified peritoneal macrophages as a prominent infected cell type at early time points post-infection. However, the state of virus infection in these cells remained in question. Though MHV68 has been reported to latently infect macrophages at late times post-infection (10), MHV68 has also been reported to lytically replicate in macrophages in vitro and in vivo (8, 30). Additionally, it remained possible that the detection of LANA::βlac in peritoneal macrophages may result from phagocytosis of other lytically-infected cells rather than bona fide infection. To address this issue, we quantified the frequency of cells capable of producing virus using a limiting dilution-based assay, comparing virus production from intact and mechanically disrupted PerCs (38). In this assay, virus production from intact cells can result from either preformed, cell-associated virions or reactivation from latency. In contrast, mechanical disruption of cells prevents reactivation from latency, a process that requires viable cells, while having minimal impact on the detection of preformed virus (38). PerCs from mock, MHV68, or MHV68.LANA::βlac-infected mice were isolated at 3 dpi and plated in a series of dilutions on a mouse embryonic fibroblast monolayer (MEFs), a sensitive and permissive indicator for virus infection. Cocultures were observed for virus-induced cytopathic effect (CPE) on MEF monolayers, quantifying preformed virus in mechanical disrupted PerCs and the sum of preformed virus and reactivation from latency in intact PerC co-cultures.

PerCs isolated from MHV68- or MHV-68.LANA::βlac infected mice demonstrated virus-induced CPE in both intact and mechanically-disrupted cells (Figure 8). In contrast, mock-infected PerCs demonstrated no detectable CPE in either condition (Figure 8). When we quantified the frequency of intact cells capable of producing virus, we found that ~1 in 7 PerCs (~14%) were associated with virus production in both MHV68 and MHV-68.LANA::βlac-infected mice. In contrast, limiting dilution analysis of mechanically-disrupted cells identified a frequency of ~1 in 100 (1%) of cells containing preformed virus (Figure 8). These data indicate that PerCs from MHV68-infected mice contain a mixture of latent and preformed virus, with latent infection comprising the dominant fraction of infected cells.

**Figure 8:**
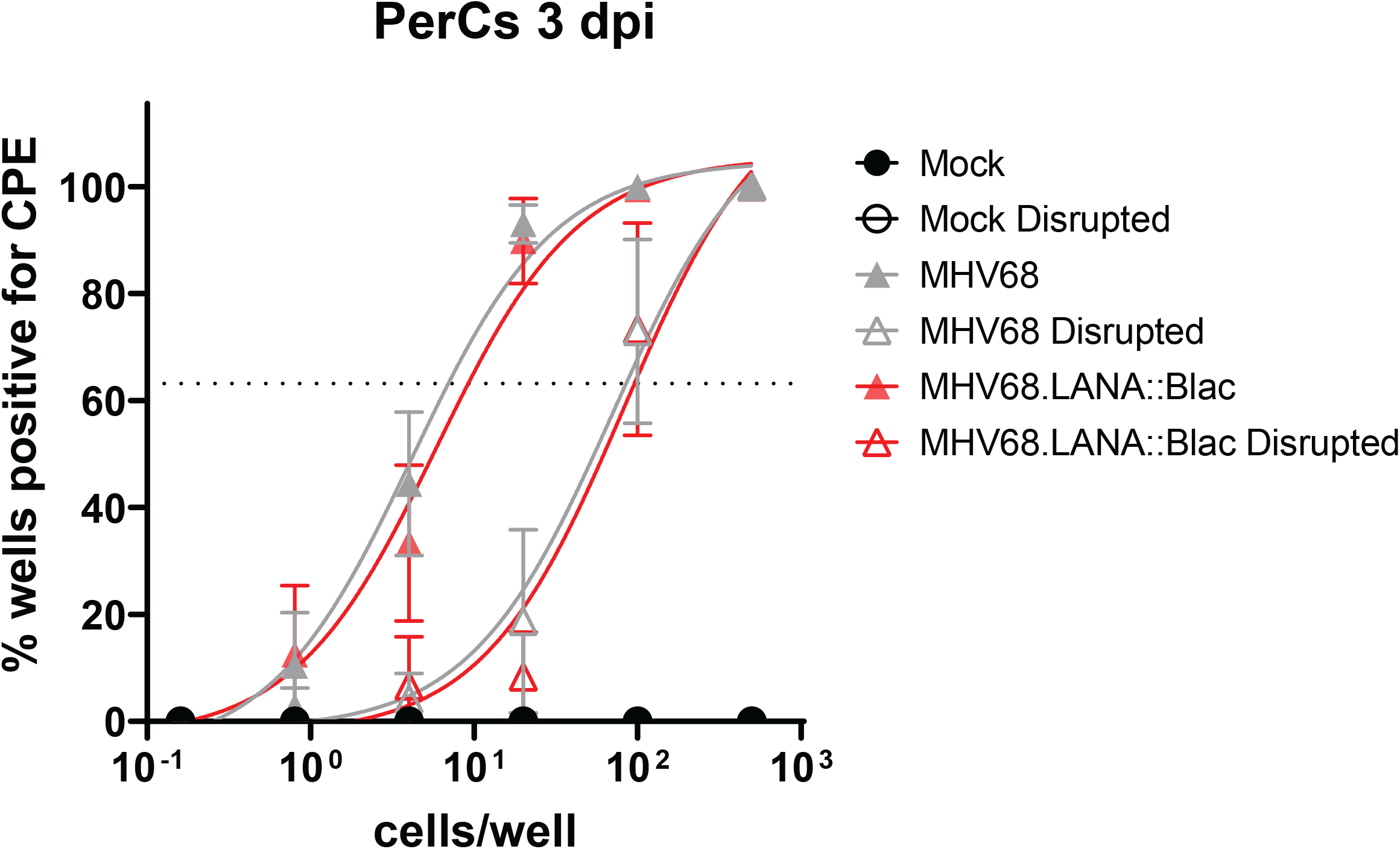
The MHV68-infected peritoneal cavity contains both lytic and latent infection at 3 days post infection. Analysis of virus infection in peritoneal cells obtained from mock, MHV68 or MHV68.LANA::βlac infected mice. PerCs were harvested at 3 dpi and subjected to limiting dilution analysis on mouse embryonic fibroblasts, with virus production quantified by cytopathic effect (CPE) 21 days post-plating. Analysis quantifies the mean frequency of wells which were positive for virus in intact and mechanically disrupted samples. 24-36 wells per dilution per condition were analyzed for undisrupted samples, and 12 wells per dilution per condition for disrupted samples. Dashed line at 63.2% is used to determine the frequency of virus-positive cells, defined by Poisson distribution. Analysis performed via nonlinear regression analysis in GraphPad Prism. Data representative of three experiments with 2 mice per condition per experiment.

## Discussion

MHV68 is an established and efficient model virus for studying both pathogen and host elements of gammaherpesvirus infection. Though much progress has been made in characterizing the virus and the response it induces, its range of target cells, its dynamic viral lifecycle, and its low frequency of infected cells during latency make it difficult to track infection at the single-cell level. The MHV68.LANA::βlac virus overcomes many of these issues by providing a highly sensitive, efficient and robust readout of infected cells detected by flow cytometric analysis, allowing the rapid analysis of thousands of cells.

While the MHV68.LANA::βlac system has been previously used to identify infected cells (12, 24, 25, 32), the broad emission spectrum of uncleaved and cleaved CCF2 has limited its use in multiparameter flow cytometry. Based on this limitation, we sought to adapt its detection to full spectrum, spectral flow cytometry, a technology that bypasses restrictions of overlapping fluorescence emission spectra through the detection of unique emission profiles, to allow for multiparameter single-cell immunophenotyping. Upon optimization of CCF2-AM substrate and incubation conditions, we were able to measure CCF2-AM fluorescence in concert with a panel of twelve surface markers, a major advance in utility of this reporter system. Notably, while spectral flow cytometry affords unique opportunities to detect the CCF2 compound, an equally important modification in this manuscript was the use of a much lower concentration of CCF2-AM substrate, circumventing the need to reduce laser voltages observed with higher CCF2-AM concentrations. Though this approach allowed detection of cleaved CCF2 fluorescence, it is of note that virtually all cells positive for cleaved CCF2 fluorescence also retain an uncleaved CCF2 fluorescence spectrum, suggesting that beta-lactamase expressing cells are unable to cleave every molecule of CCF2-AM within the cytoplasm. It is unclear whether there is an ideal concentration of CCF2-AM substrate that would allow for complete cleavage by beta-lactamase, and whether this would be desirable, given that detection of the uncleaved substrate also provides valuable evidence of equal substrate loading. Following identification of these optimized conditions, we further established an approach to ensure full spectrum unmixing by using cells with a high cleaved CCF2 fluorescence for spectral unmixing.

With the optimization of MHV68.LANA::βlac detection by spectral flow cytometry, we characterized the impact of acute MHV68 infection on cells in the peritoneal cavity, examining both the broad impact of MHV68 infection as well as the specific targets of MHV68 infection. We found that MHV68 infection is associated with pronounced changes in the cellular composition of PerCs, with rapid increases in the frequency and number of CD8 T cells, and pronounced changes to myeloid cells, including transient elimination of large peritoneal macrophages coupled with expansion of small peritoneal macrophages (SPMs) and monocytes. MHV68 infection was further associated with long-term phenotypic changes in LPMs and SPMs, including upregulation of MHC II expression in LPMs.

One of the most pronounced changes in the peritoneal cavity was the transient depletion of LPMs. The disappearance of LPMs, a phenomenon referred to as the ‘macrophage disappearance reaction’, is a well-known outcome in response to inflammatory stimuli, including thioglycollate, LPS, virus and parasitic infection (29, 37, 39). During this response, LPMs are presumed to traffic to the omentum, where they potentially carry antigen to stimulate B cells to mature in germinal centers present in milky spots, small lymphoid structures within the omentum (40). As the omentum is a highly vascularized tissue, it is also possible that macrophages further disseminate to the lymphatics or the bloodstream (41). We did not see any sign of death in LPMs before elimination, nor did we observe their presence when adding additional EDTA to our extraction solution (data not shown), suggesting that the loss of LPMs is not due to their lysis or adherence to the peritoneal walls.

Despite the depletion of LPMs at 8 dpi, LPM-like cells reappeared in the infected peritoneal cavity by 14 dpi. It is unclear how this subset is repopulated, as LPMs were originally described as tissue-resident and self-renewing (29). One alternate explanation could be that LPMs are replaced by SPMs or monocytes, as these cell subsets exhibit similar F4/80 and CD11b expression during the recovery phase, with LPM and SPM characterized by relatively comparable expression of MHC II and CD86 at later times post-infection. The potential replacement of LPMs by SPMs or monocytes is supported by previous fate-mapping studies (42, 43) and represents an important unresolved question in the context of MHV68 infection.

In addition to observing widespread changes in peritoneal cell composition, our studies allowed detailed analysis of targets of early MHV68 infection in the peritoneal cavity. This analysis revealed a high frequency of infection (25-40%) in large peritoneal macrophages through 3 dpi. At 8 dpi, the SPM/monocyte population becomes the predominant source of infection. We note that the majority of LANA::βlac+ within the SPM/monocyte population express high levels of MHCII, suggesting these cells are SPMs rather than monocytes (29). Dendritic cells and B cells represented a minor fraction of LANA::βlac+ events. We further identified that virus infection resulted in changes to macrophage phenotype, both within virus-infected cells and in uninfected cells. Among these, an increased frequency of LANA::βlac+ LPMs were characterized by high expression of MHC II expression, suggesting that these cells may either be activated and/or directly capable of presenting antigen to virus-specific CD4 T cells. The observation that MHV68 is predominant in LPMs before their disappearance at 8 dpi also suggests that MHV68 may potentially exploit LPM responses to inflammation to facilitate virus dissemination to the omentum. Consistent with this, previous studies by Gray et al identified MHV68+ cells in the omentum, including B220+ B cells and a smaller fraction of CD11b+ cells, presumed to be macrophages (11). It is also possible that MHV68+ LPMs either undergo lytic replication and/or are directly targeted for destruction by antiviral T cells.

Although analysis of virus infection was done on bulk PerCs at 3 dpi, a time when LPMs were the dominant MHV68+ population, it is notable that PerCs contained a mixture of latently infected cells and cells undergoing lytic replication. While the definition of latency is often precluded by the presence of lytic infection, the comparable frequency of LANA::βlac+ cells, defined by flow cytometry, and the frequency of cells capable of reactivating from latency, defined by limiting dilution analysis, is consistent with latent infection in the majority of LPMs. These data strongly suggest that MHV68 latent infection in PerCs can be found as early as 3 dpi, a result that is consistent with a previous report identifying the early establishment of latency in the lung following intranasal infection (44). While this analysis focused on bulk PerCs, future studies purifying LANA::βlac+ peritoneal macrophages may provide further evidence for latent infection during these early stages of infection.

We note that at 14 dpi, the LANA:: βlac is not well detected above background. Previous studies using this reporter virus reported LANA::βlac+ cells at less than or equal to 0.1% of splenocytes at 16 dpi (24, 45), and limiting dilution qPCR experiments have reported frequencies of less than 1% infected cells in specific cell populations in the peritoneal cavity at 16 dpi (17). Based on these data, we infer LANA:: βlac+ cells are a rare but present population within the peritoneal cavity at 14 days or later post infection. These cells may be more effectively detected by greatly increasing the number of peritoneal cells analyzed and/or by enriching for cell types known to be infected prior to analysis.

In total, this study demonstrates optimization of the MHV68.LANA::βlac reporter system, to identify targets of virus infection using spectral flow cytometry. By tracking MHV68 infection in the peritoneal cavity, this analysis identified both broad changes in the immune landscape of the peritoneal cavity as well as differential protein expression profiles in virus-infected and uninfected cells. These studies identify MHV68-induced myeloid reprogramming, characterized by the induction of MHC II in large peritoneal macrophages and CD86 induction in small peritoneal macrophages, consistent with virus-induced alterations to the peritoneal cavity previously associated with MHV68-induced cross-protection against bacterial infection (46). We further identify peritoneal macrophages as a prominent target of acute infection and identify a mixture of lytic and latent infection early post-infection. This work emphasizes the dynamic interactions between MHV68 and the host, and provide critical cellular context that will allow future analysis of factors that regulate early virus-host dynamics.

## Acknowledgements

We thank the University of Colorado Anschutz School of Medicine Office of Laboratory Animal Resources (OLAR) and Alexander Sosa for technical support with animal husbandry. We would also like to thank the University of Colorado Cancer Center Flow Cytometry Shared Resource as well as the Department of Immunology & Microbiology Flow Cytometry Shared Resource Lab for assistance with flow cytometry. Funding for this work was provided by NIH grant AI32419 (LJB).The Flow Cytometry Shared Resources of the University of Colorado Cancer Center receive direct funding support from the National Cancer Institute through Cancer Center Support Grant P30CA046934.

---

**Supplementary Figure 1:**
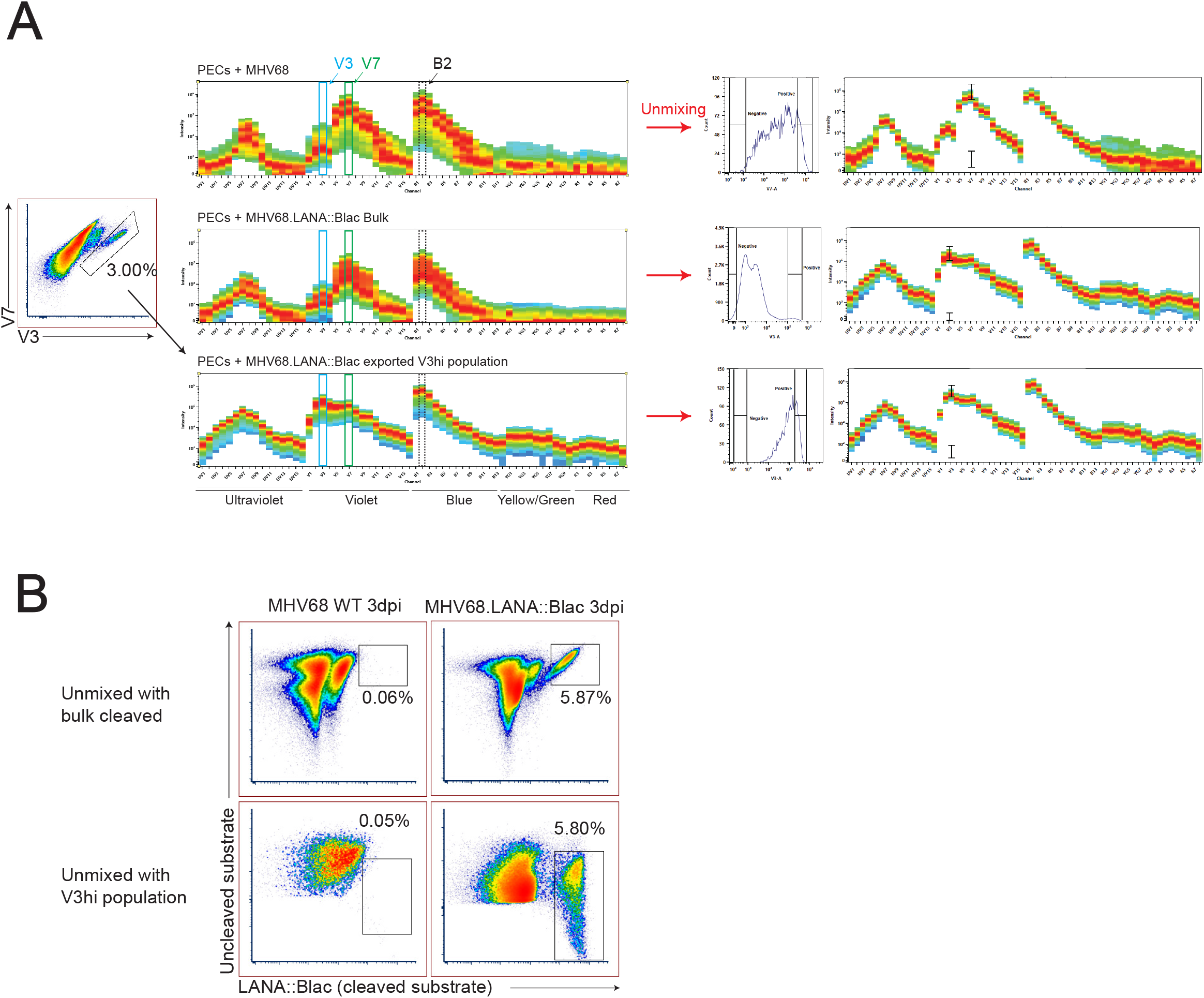
Strategy for spectral unmixing with CCF2 on the spectral flow cytometer when the frequency of cleaved CCF2 events is low. (A) Raw spectral data from a five laser Cytek Aurora spectral flow cytometer from samples at 3 dpi, analyzing fluorescence in peritoneal cells obtained from MHV68 or MHV68.LANA::βlac infected mice. Emission in V3, V7 and B2 detectors is highlighted. Top panel: uncleaved CCF2-AM signal remains robust, regardless of time point post infection, allowing for easy creation of a single stain control. Middle panel, right: when cleaved CCF2-AM signal is hard to discern, a new single stain control can be created by plotting emission in the V3 and V7 detectors (left) and gating on cells that have high emission in the V3 detector. Bottom panel: This population can be exported as a new FCS file and then reimported back into the experiment, resulting in a more robust single stain control. (B) Unmixed data comparing the two unmixing strategies for cleaved CCF2. Data utilized is from peritoneal exudate cells of mice infected with MHV68 or MHV68.LANA::βlac for three days. Unmixing with bulk cells or with the exported V3^High^ population produces a similar percentage of cells positive for cleaved CCF2.

**Supplementary Figure 2:**
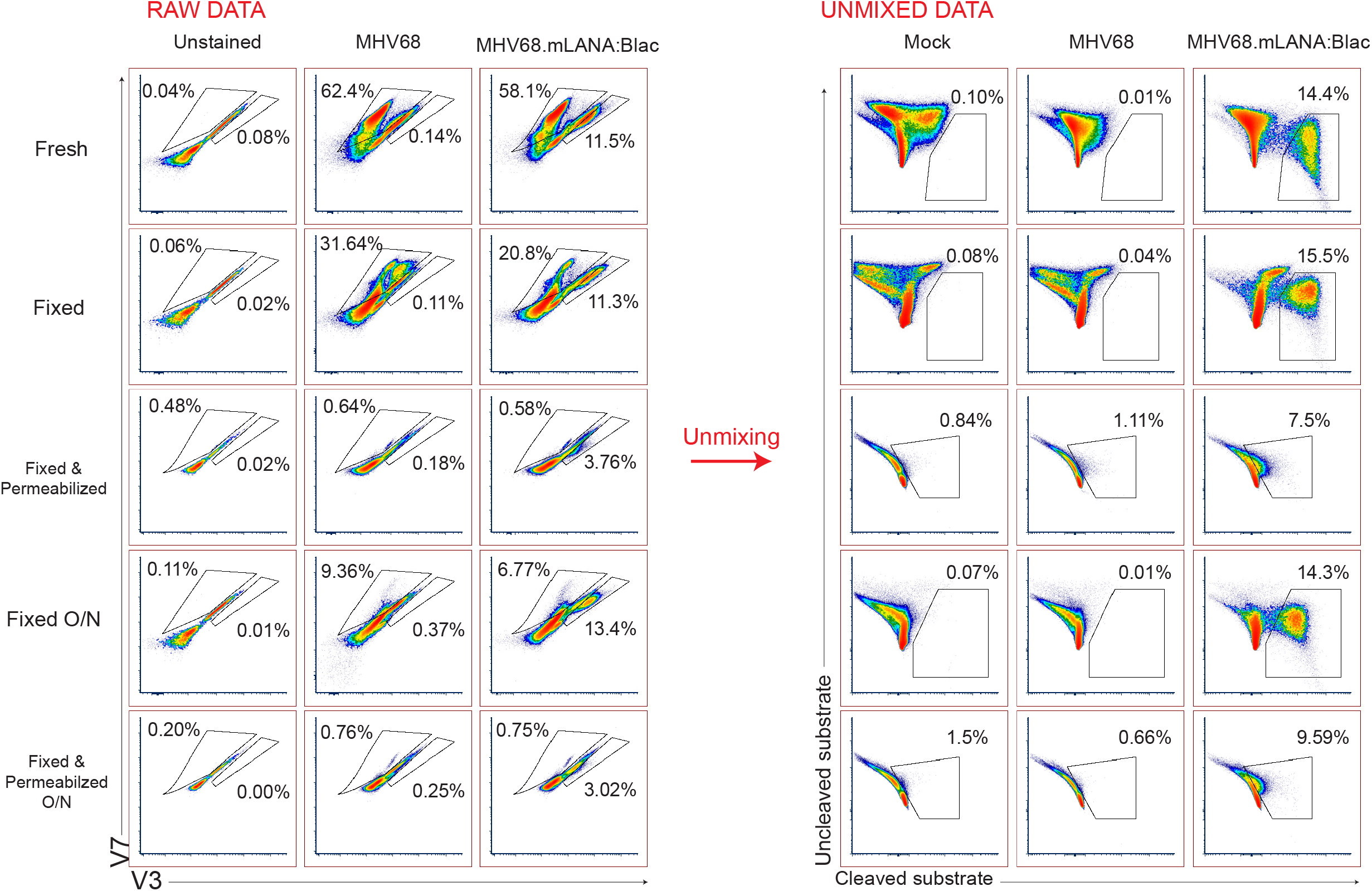
The impact of fixation and permeabilization on CCF2-AM detection by spectral flow cytometry. PerCs were extracted from mice and infected *in vitro* with WT MHV68 or MHV68.LANA::βlac. At 20 hours post-infection, cells were harvested and either left unstained or stained with CCF2-AM and subjected to one of the following conditions: (1) Run “Fresh” on the spectral flow cytometer the day of harvest, (2) Fixation using 4% paraformaldehyde and run immediately after fixation (“Fixed”), (3) Fixation and Permeabilization and run immediately after (“Fixed and Permeabilized”), (4) Overnight fixation and run the following day (“Fixed O/N”), or (5) Overnight fixation and permeabilization and run the following day (“Fixed & Permeabilized O/N”). Left column depicts raw data, showing fluorescence in the V3 and V7 detectors, with right column depicting unmixed data for cleaved and uncleaved CCF2 fluorescence. Gating was drawn based on unstained controls, and samples were gated on live singlets. Unmixing was performed using controls that underwent the same staining conditions or, in the case of insufficient signal, with controls from other conditions with robust signal. Gating for unmixed data was drawn based on mock and MHV68 WT-infected controls. Data representative of two experiments.

**Supplementary Figure 3:**
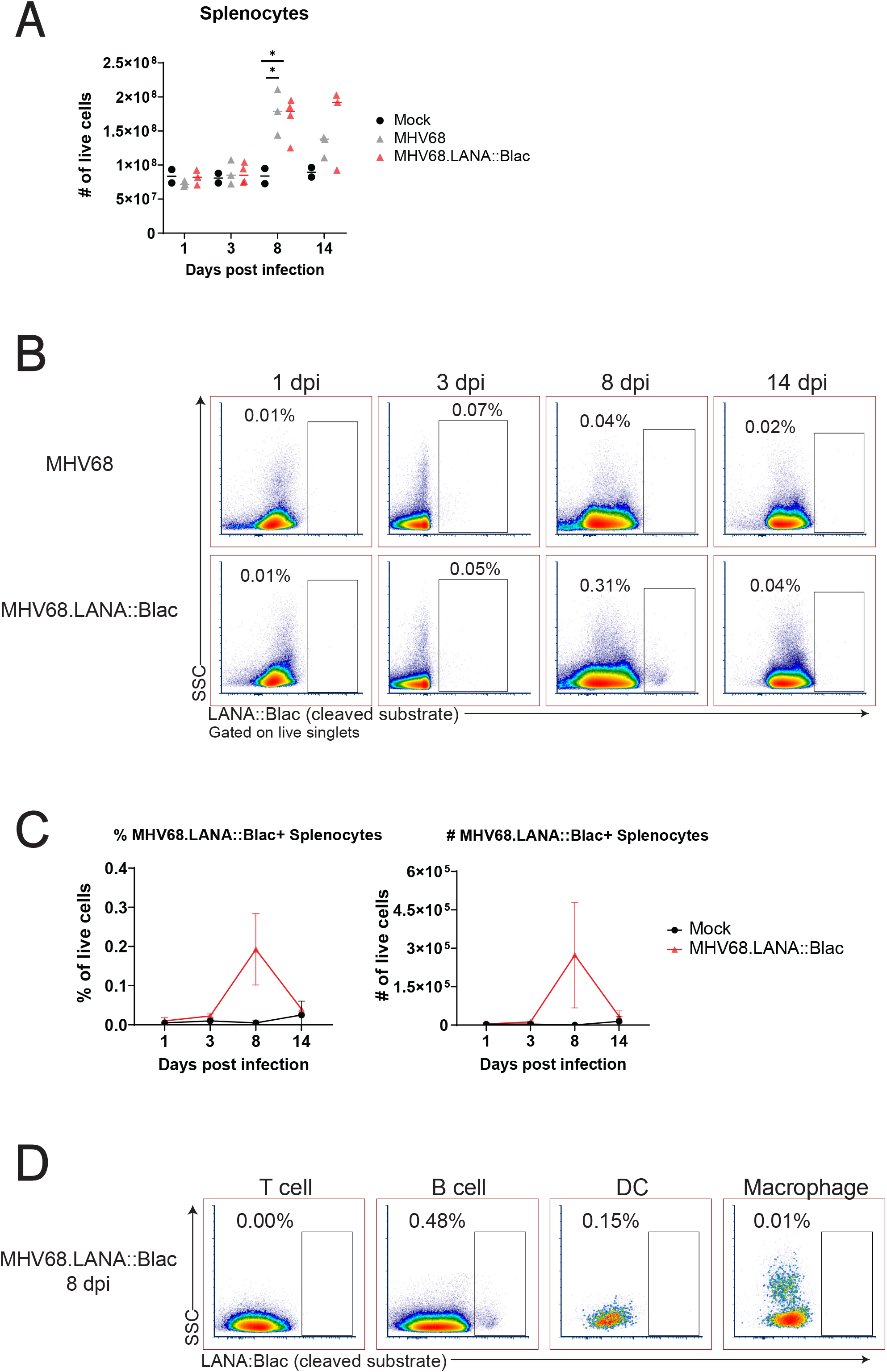
MHV68.LANA::βlac is detectable in B cells at 8 days post infection in the spleen. C57BL/6 mice were infected i.p. with 1 million PFUs of WT MHV68 or MHV68.LANA::βlac. At 1, 3, 8 and 14 dpi, mice were sacrificed and spleens were harvested, counted, and stained with CCF2-AM, viability, and surface markers. (A) Viable cell counts of harvested spleens in mock, MHV68 or MHV68.LANA::βlac infected mice. (B) Dot plots showing the frequency of splenocytes positive for cleaved CCF2. LANA::βlac signal is enriched above background (defined by WT MHV68 infection) at 8 dpi. (C) Quantification of the mean frequency and number of LANA::βlac+ cells shown in (B). (D) Dot plots showing percentage of T cells, B cells, dendritic cells and macrophages that are positive for cleaved CCF2. B cells are the only population with cleaved CCF2 signal, with about 0.5% of cells staining positive. Data representative of three experiments with 2-3 mice per condition per experiment.

**Supplementary Figure 4:**
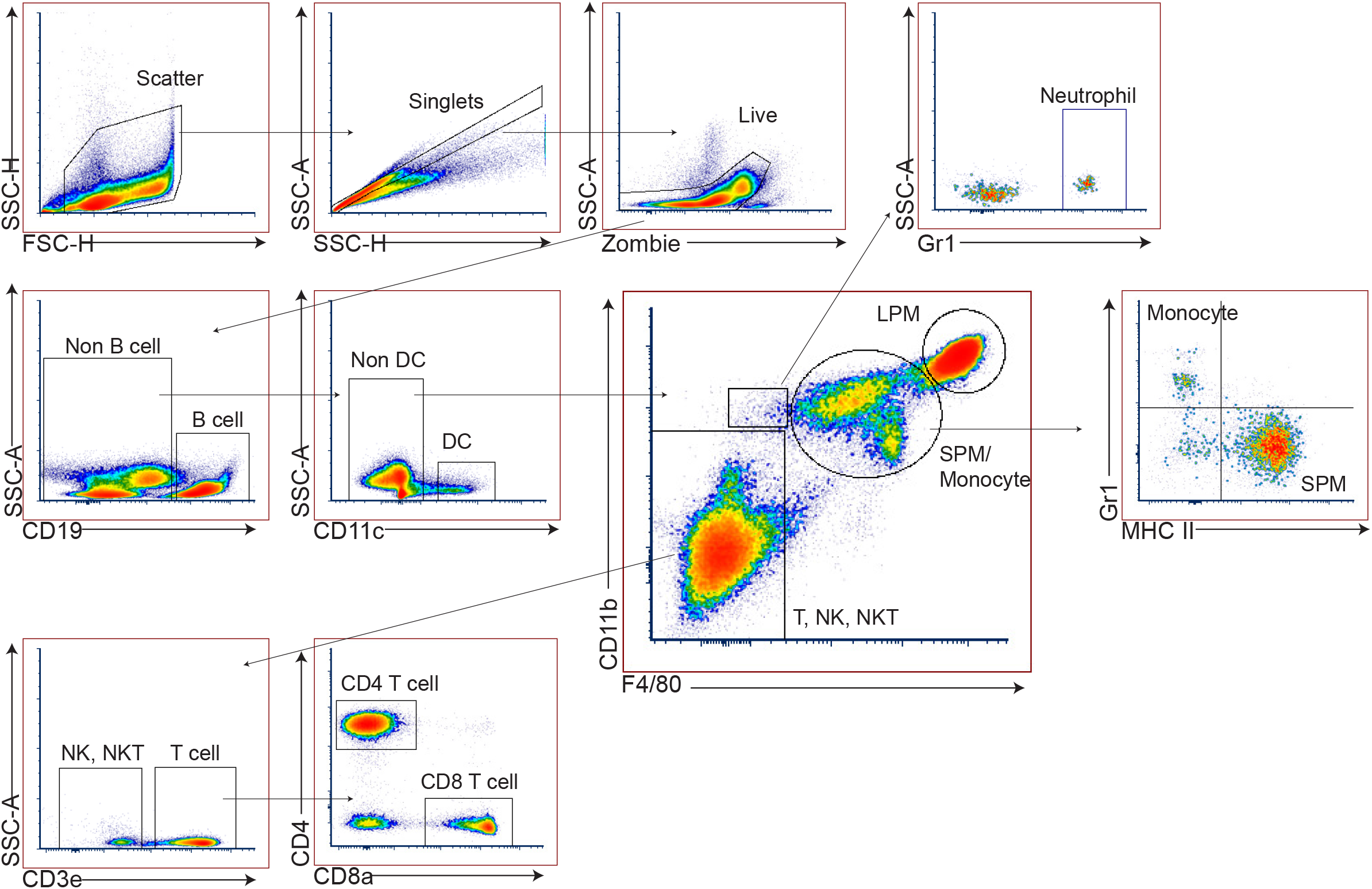
Gating strategy for analysis of spectral flow cytometry data. Representative gating scheme adopted from *Ghosn et al 2010.* Sample derived from PerCs of a mock-infected mouse.

